# Claustrum volume in humans – lifespan trajectory and effect of age, hemisphere, and sex

**DOI:** 10.1101/2025.09.09.675215

**Authors:** Sevilay Ayyildiz, Antonia Neubauer, Melissa Thalhammer, Hongwei Bran Li, Jil Wendt, Aurore Menegaux, Rebecca Hippen, Benita Schmitz-Koep, David Schinz, Claus Zimmer, Behcet Ayyildiz, Abdullah Ors, Belgin Bamac, Dennis M. Hedderich, Christian Sorg

## Abstract

The human claustrum is a bilateral, thin, irregularly shaped gray matter structure located between the striatum and insula. While previous research demonstrated the effect of distinct medical conditions, such as prematurity, schizophrenia, and Alzheimer’s disease, on claustrum function and structure, it is poorly understood how non-pathologic biological conditions effect the claustrum. This study aimed to investigate the effect of age, hemisphere, and sex on claustrum volume.

We used T1-weighted 3 Tesla MRI scans of 3,474 healthy participants ranging from 1 to 80 years of age, deep learning-based automated claustrum segmentation, and a normative modeling approach to delineate lifespan trajectories of claustrum volumes for both hemispheres and sexes. Additionally, ordinary least squares regression analyses were applied to further characterize age, hemisphere, and sex effect.

Lifespan analysis revealed a trajectory of rapid claustrum volume increase from infancy to adolescence (∼ 1–15 years, annual growth 39.300 mm^3^/year), a plateau phase from early to middle adulthood (∼ 15–40 years, annual change 0.153 mm^3^/year), and a subsequent decline from middle adulthood to old age (∼ 40–80 years, annual decrease 10.325 mm^3^/year). The right claustrum was on average larger than the left one across all ages. Finally, overall, females had larger total intracranial volume-adjusted claustrum volumes than males across the lifespan.

Results demonstrate a distinct effect of age, hemisphere, and sex on claustrum volume. Data provide a comprehensive framework for sex- and hemisphere-sensitive claustrum structure lifespan trajectories relevant for studying neurodevelopmental and neurodegenerative effects on the claustrum.

## 1. Introduction

Nestled deep within the human brain, the claustrum is a small grey matter structure located between the external and extreme capsule, below the insular cortex and nearby the striatum (Crick & Koch, 2005; Jackson, Smith, & Lee, 2020). In mammals, in general, it originates from primordial cells in radial progenitor domains of lateral pallium derivatives early during brain development (Bruguier et al., 2020). There is a dynamic interplay between radial and tangential migration processes, which contribute to claustrum formation. Furthermore, transient cell populations, namely subplate neurons and preoligodendrocytes, are also involved in claustrum development (Watson and Puelles, 2017; Smith et al., 2019; Bruguier et al., 2020). Relative to its volume, the claustrum is one of the most extensively connected brain structure in the brain and builds networks with cortical and subcortical regions (McBride et al., 2023; Torgerson, Irimia, Goh, & Van Horn, 2015; Wendt et al., 2024). It establishes regionally distinct reciprocal microcircuits with cortical layers, exerting selective inhibitory control over cortical columns, thus modulating a wide range of processes such as attentional processes (Goll, Atlan, & Citri, 2015; White et al., 2020), salience detection (Smith et al., 2019), cognitive control (Krimmel et al., 2019; Madden et al., 2022), task switching (Huang et al., 2024), behavioral engagement (Atlan et al., 2021; J. Liu et al., 2019), pain processing (Stewart et al., 2024), and sleep regulation (Atlan et al., 2024; Narikiyo et al., 2020).

The developmental and aging trajectories of the claustrum are vulnerable to disruption, leading to macro- and microstructural deviations in varying brain diseases with putative functional correlates (Bruen, McGeown, Shanks, & Venneri, 2008; Cascella et al, 2011a; Davis, 2008; Wang et al., 2023). For example, premature birth affects perinatal claustrum development, followed by altered claustrum macro- and microstructure in newborns (Neubauer et al., 2023), with these changes partly lasting into adulthood and being related to general cognitive performance (Hedderich et al., 2021). In patients with schizophrenia, a further example of a neurodevelopmental disorder, claustrum volumes are more than 10% lower compared to healthy controls, with these aberrations being correlated with attentional deficits (Bernstein et al., 2016; Schinz et al., 2024). Finally, concerning aging and aging-related neurodegenerative disorders, decreased claustrum volumes have been observed in Alzheimer’s (Bruen et al., 2008; Ayyildiz et al., 2023) and Parkinson’s disease (Shao, Yang, & Shang, 2015). While these human functional and clinical studies of sex-mixed samples of different ages clearly demonstrate the effect of certain functional or clinical conditions on claustrum features, the effect of more basic biological conditions such as age, hemisphere, or sex is unknown. In order to not only better understand claustrum development but also to provide a framework to interpret the above-mentioned influences of neurodevelopmental, neurodegenerative, and functional conditions, we investigated the effect of age, sex, and hemisphere on claustrum volume across the lifespan.

To investigate these factors, we focused on claustrum volumes as measured by T1-weighted MRI and subsequent automated machine learning-based claustrum segmentation. Human claustrum studies are rare due to the complex, thin, and irregular claustrum shape that makes claustrum segmentation challenging. Furthermore, large sample studies on the claustrum are even rarer due to the technical difficulties of large-scale and reliable claustrum segmentation. Recently, an in-house solution to this challenge was proposed using deep learning-based segmentation approaches in 3 Tesla anatomical MRI, enabling reliable claustrum volumetry in larger samples of hundreds of adult and neonate subjects (Li et al., 2021a; Neubauer et al., 2022a; Schinz et al., 2024; Wendt et al., 2024). Here, we applied this approach to several thousands of healthy subjects, from 1 year old to older adults of 80 years, derived from multiple public MRI data. Using a sample of 3,474 healthy participants across 24 sites, we performed normative modeling to estimate lifespan trajectories for averaged claustrum volume as well as left and right claustrum volumes of females and males, respectively. Using large datasets, normative modeling methods can be applied to accurately model brain development, spanning the human lifespan by taking into account not only the mean but also the variance of the population for a certain brain measure. To date, normative modeling has primarily been used on cortical features such as thickness or subcortical volumes, excluding the claustrum (Bethlehem et al., 2022a; Chen, Holmes, Zuo, & Dong, 2021; Rutherford et al., 2022). Additionally, we compared these sex- and hemisphere-distinct trajectories using a canonical ordinary least squares (OLS) approach to test for more specific age, hemisphere, and sex influences on claustrum volumes.

## 2 Materials and Methods

### 2.1. Datasets and participants

The study includes data from six datasets, comprising 3,474 healthy participants from 24 sites, ranging from 1 to 80 years of age, with nearly equal representation of men and women (48% and 52%, respectively) (Figure 1, Table 1). The datasets include anonymized conventional 3D T1-weighted (T1w) 3 Tesla MRIs obtained from the following sources: the UNC/UMN Lifespan Baby Connectome Project (BCP), the Calgary Preschool MRI Dataset (CPD) (https://osf.io/axz5r/), a selected dataset from the Adolescent Brain Cognitive Development (ABCD) study, 5.1 release, primarily including participants aged between 9 and 12 years (https://abcdstudy.org), the Human Connectome Project (HCP) Development dataset, the HCP Young Adult dataset, and the HCP Aging dataset (https://www.humanconnectome.org). All participants underwent screening to ensure the absence of mental disorders, cognitive impairment, or significant medical morbidity, including brain abnormalities (Howell et al., 2019; Reynolds, Long, Paniukov, Bagshawe, & Lebel, 2020; Somerville et al., 2018; Van Essen, Smith, Barch, Behrens, & Yacoub, 2013; Volkow et al., 2018).

**Table 1:**
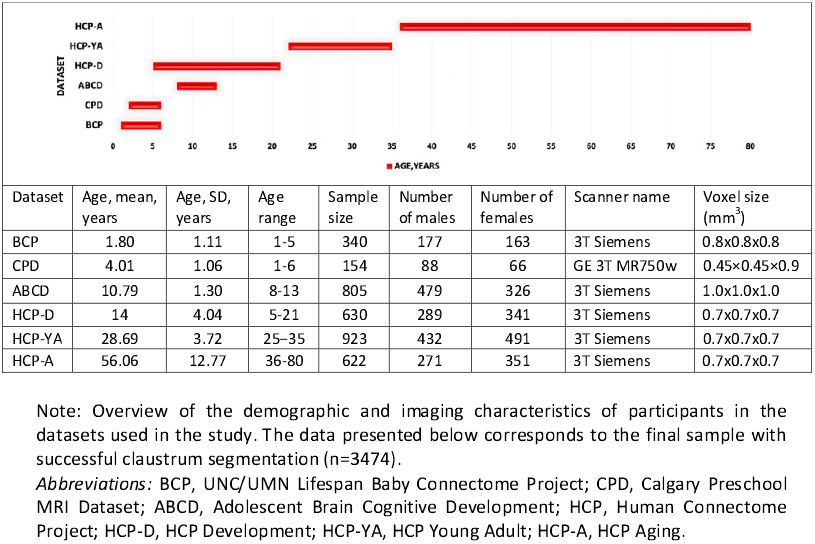
Demographic and imaging characteristics across different data sets.

**Figure 1:**
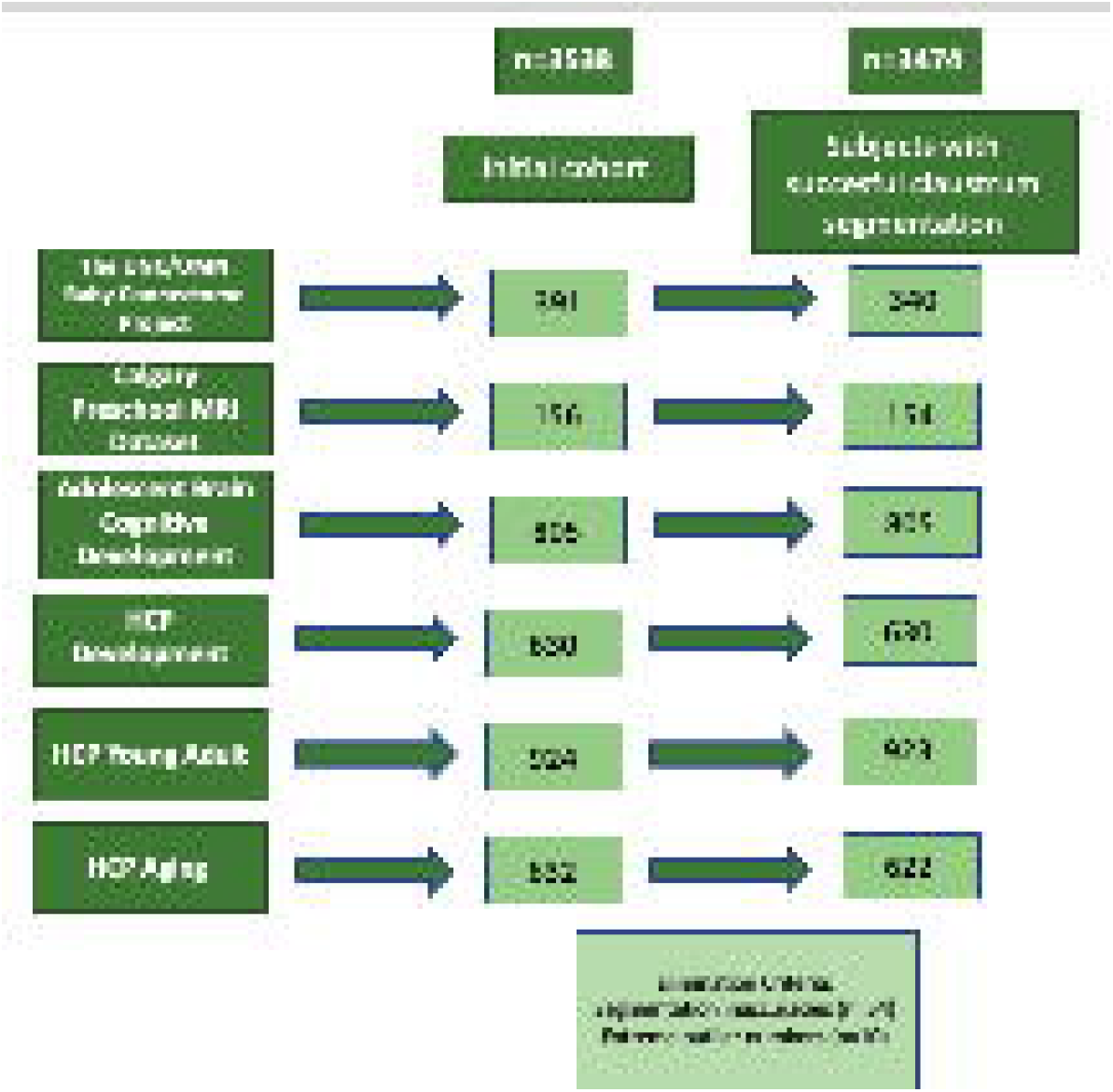
Flow diagram of participants included in the study. The diagram illustrates the initial cohorts of participants from six datasets, detailing the number of subjects included in the study after exclusion due to segmentation inaccuracies and extreme outlier numbers in claustrum volume. *Abbreviations:* HCP, human connectome project

### 2.2. Imaging acquisition

Detailed information on scanner vendors, magnet strengths, and acquisition parameters for each sample are shown in Table 2. Basic quality control of all brain MR images was conducted by researchers of the respective dataset (see next for details) (Howell et al., 2019; Reynolds, Long, Paniukov, Bagshawe, & Lebel, 2020; Somerville et al., 2018; Van Essen, Smith, Barch, Behrens, & Yacoub, 2013; Volkow et al., 2018). The study only included participants whose T1w MRI scans had enough resolution to segment the claustrum accurately (i.e., spatial voxel resolution 1x1x1mm^3^ or lower, based on previous studies (Li et al., 2021a; Neubauer et al., 2022a).

**Table 2:** Image acquisition parameters of structural image and scanner details for datasets used in the study.

| Dataset | Age range of dataset | Scanner details | Imaging parameters | Reference (screening process and eligibility criteria) |
| --- | --- | --- | --- | --- |
| **Lifespan Baby Connectome Project (BCP)** | 0-60 months | 3T Siemens scanner, 32-channel head coil, FOV: 256×256, Matrix: 320×320, 208 sagittal slices | 0.8 mm<sup>3</sup> isotropic resolution, TR = 2400 ms, TE = 2.24 ms, FA = 8° | (Howell et al., 2019) |
| **The Calgary Preschool MRI Dataset (CPD)** | 2-8 years | GE 3T MR750w system, 32-channel head coil, FOV: 256×256, Matrix: 512×512, 210 sagittal slices | 0.9 x 0.9 x 0.9 mm<sup>3</sup> isotropic resolution, TR = 8.23 ms, TE = 3.76 ms, FA = 12°, Inversion Time = 540 ms | (Reynolds, Long, Paniukov, Bagshawe, & Lebel, 2020) |
| **Adolescent Brain Cognitive Development Dataset (ABCD)** | from ages 9–10 years through adulthood | Siemens, GE, Philips MR scanners, 21 sites, FOV: 256×256, Matrix: 256×256, 176 sagittal slices | 1 mm<sup>3</sup> isotropic resolution, TR = 2500 ms, TE = 2.88 ms, FA = 8° | Volkow et al. (2018) |
| **HCP Young Adult (HCP-YA)** | 22-37 years | 3T Siemens scanner, 32-channel head coil, FOV: 224 x 240, Matrix: 320 x 320, 256 sagittal slices | 0.7 mm<sup>3</sup> isotropic resolution, TR = 2400 ms, TE = 2.14 ms, FA = 8° | Van Essen et al. (2013) |
| **HCP Development (HCP-D)** | 5-21 years | 3T Siemens scanner, 32-channel head coil, FOV: 256×240×166, Matrix: 320×300×208 slices | 0.7 mm<sup>3</sup> isotropic resolution, TR = 2500 ms, TE = 1.8/3.6/5.4/7.2 ms, FA = 8° | (Somerville et al., 2018) |
| **HCP Aging (HCP-A)** | 36-100+ years | 3T Siemens scanner, 32-channel head coil, FOV: 256×240×166, Matrix: 320×300×208 slices | 0.7 mm<sup>3</sup> isotropic resolution, TR = 2500 ms, TE = 1.8/3.6/5.4/7.2 ms, FA = 8° | (Somerville et al., 2018) |

Note: Overview of the age range of participants, the scanner details, imaging parameters of T1-weighted (T1w) images, and reference details of the screening process and eligibility criteria for the datasets used in this study.
*Abbreviations:* HCP, human connectome project; TR, Repetition time; TE, Echo time; FA, Flip angle.

### 2.3. Claustrum segmentation

Briefly, we used a deep learning-based algorithm to segment the claustrum in T1w MRIs from this diverse sample of participants, aged 1-80 years, from multi-site neuroimaging data sets, using transfer learning from a pre-trained claustrum segmentation model developed for T1w scans of adults (Li et al., 2021a; Neubauer et al., 2022a).

#### 2.3.1. Data Preparation

Prior to claustrum segmentation, the following preprocessing steps were applied to the T1w MRI scans. Initially, structural preprocessing was performed using the FSL anatomical processing script (fsl_anat). The pipeline performed brain extraction and bias field correction. Subsequently, denoising was performed using the Pierrick Coupe MATLAB-based MRI Denoising Software (Coupe et al., 2008), employing the Optimized Nonlocal Means option to reduce noise while preserving image details. The images were resampled to a voxel size of 1 mm^3^ isotropic to maintain consistency across all scans using fslmaths. This resampling was imperative to meet the requirements of the subsequent claustrum segmentation step, necessitating images to be of identical resolution.

Additionally, the total intracranial volume (TIV) was computed for each subject using FSL-FAST (FMRIB’s Automated Segmentation Tool, version 6.0.2). Further steps were undertaken on top of the basic preprocessing measures to uphold consistency in dimensions and orientation across diverse subjects. This entailed cropping or padding volumes to a standardized size (200⍰ × ⍰ 200) to ensure uniform input size for the deep-learning model. Z-score normalization was also applied to MRI volumes to standardize intensity values as performed in previous studies (Li et al., 2021a; Neubauer et al., 2022a).

#### 2.3.2. Lifespan-Based Claustrum Segmentation Model: Training, Evaluation, and Application

Using T1w MRI scans from a lifespan sample and corresponding manual claustrum segmentations of a selected sub-sample by an experienced neuroanatomist (S.A.), we trained and aggregated three axial and coronal view claustrum models into a unified claustrum model at the voxel level. The details of the training model process can be found in the Supplementary material. In brief, three axial and three coronal models were trained for 30 epochs, using a batch size of 60 and a learning rate of 0.0002, on 40 subjects selected from across all datasets, ensuring balanced representation across various age groups and sites/scanners. Data augmentation techniques were applied to improve model robustness, including scaling, rotation, shifting, shearing, and intensity modification. Then, to enhance model robustness further, the three axial and coronal view models were aggregated in a unified model at the voxel level.

To evaluate the model’s segmentation performance, we calculated the volumetric similarity (VS), Hausdorff distance (95th percentile) (HD95), and Dice similarity coefficient (DSC) metrics comparing the model’s output and ground truth manual segmentation in a separate test set of 10 scans, as delineated in prior studies (Li et al., 2021b; Neubauer et al., 2022b).Eventually, the resulting combined model was applied to the rest of the participants (Figure 2).

**Figure 2:**
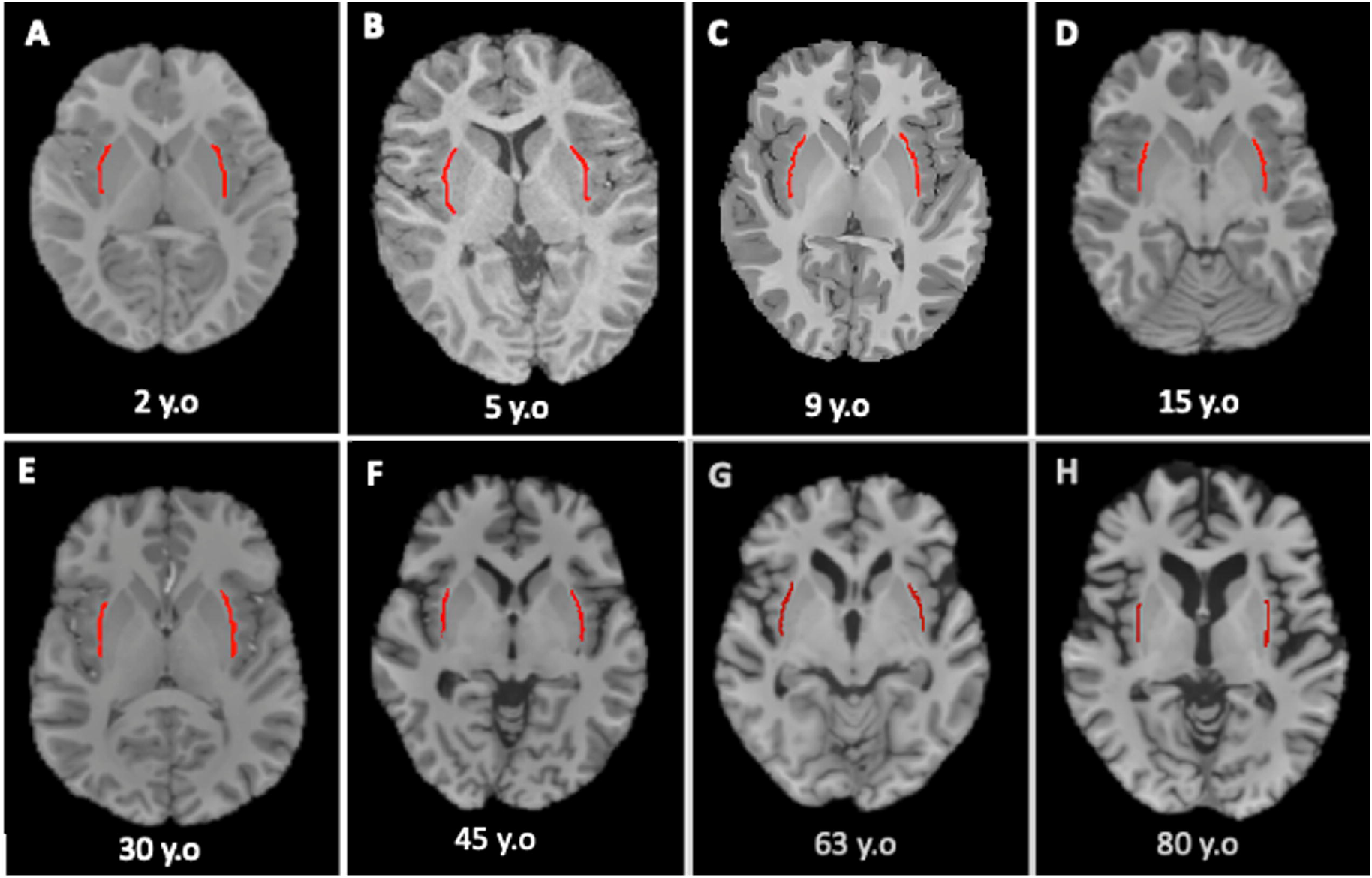
Examples of automated claustrum segmentations across the lifespan. The segmentation results were generated using an automated deep learning algorithm for T1-weighted MRI scans. *Abbreviations:* y.o, years old

#### 2.3.3. Segmentation quality assessment across data sets

After applying a deep learning algorithm for claustrum segmentation across the whole T1w MRI data set, statistical outliers of claustrum volume were identified separately within each single-year age group. Outliers were defined as instances falling below Q1 - 1.5 * IQR or above Q3 + 1.5 * IQR for each claustrum (Tukey, 1977). The segmentation quality of these outliers was meticulously scrutinized using ITK-SNAP-v 4.0.2 (Yushkevich et al., 2006). Subsequently, 64 subjects were excluded from further analyses due to segmentation inaccuracies (i.e., poor image quality, insufficient landmark visualization, poor contrast with adjacent structures) (n=54) and extreme outlier values in claustrum volume (n=10) (Figure1).

### 2.4. Statistical Analysis

#### 2.4.1. Normative modelling of averaged claustrum volume lifespan trajectories

We used the Predictive Clinical Neuroscience toolkit (PCNtoolkit) (https://pcntoolkit.readthedocs.io/en/latest) framework to model normative trajectories for averaged, left and right claustrum volume across the lifespan. A Bayesian Linear Regression model (BLR; Fraza, Dinga, Beckmann, & Marquand, 2021) with likelihood warping was used to predict the claustrum volume from a vector of covariates including sex, age, and TIV across a large healthy sample, referred to as a training set. A B-spline basis expansion was applied to the covariate vector to allow flexible modeling of non-linear effects. Cubic splines with three evenly spaced knots were selected to balance model complexity and the ability to fit covariate-related changes. Non-Gaussianity was modeled with a likelihood warping approach (Fraza, Dinga, Beckmann, & Marquand, 2021) using a “sinarcsinsh” warping function. Site effects were modeled using fixed effects, which was previously shown to effectively remove variance attributable to the scanning site (Kia et al. 2021, Rutherford et al. 2022). Hyperparameter optimization was controlled by minimizing the negative log-likelihood and by a fast numerical optimization algorithm (L-BFGS). An 80/20 train/test split stratified by site was performed to train the model on 80 % of the data and assess generalizability of the model to unseen data on the remaining 20 %. Deviation scores (Z-scores) are calculated for the n-th subject and d-th brain area as:

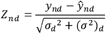

Where *yn*_*d*_ is the true outcome measure (i.e., claustrum volume), *ŷ*_*nd*_ is the predicted outcome measure, *σ*_*d*_^*2*^ is the estimated noise variance (reflecting uncertainty in the data), and (*σ*^*2*^)_*d*_ is the variance attributed to the modeling uncertainty (Rutherford 2022, Marquand 2016). Diverse error metrics, including Negative Log Likelihood (NLL), Explained Variance (EV), Mean Standardized Log Loss (MSLL), Bayesian Information Criteria (BIC), and skew and kurtosis of the Z-distribution, were computed on the test set to estimate the efficacy of the model. Visual representations were generated to evaluate test set predictions and estimate uncertainty across distinct demographic strata and age spans (Rutherford et al., 2021).

After modeling the trajectories of averaged claustrum volume, the output from the BLR model (50th percentile) was used to calculate the rate of change in claustrum volume with respect to age. We applied a piecewise linear approximation to age-related volume changes, highlighting critical phases of growth and decline in averaged claustrum volume. Within each segment, i.e., increase phase (1–15 years), plateau phase (15–40 years), and decrease phase (40–80 years), we computed linear slopes using least squares regression, representing the rate of change in claustrum volume across age.

#### 2.4.2. Hemisphere and sex differences in claustrum volume across the lifespan

##### Lifespan normative trajectory of sex- and hemisphere stratified claustrum volumes

After normatively model averaged claustrum volume lifespan trajectories, further models of sex- and hemisphere stratified claustrum volumes were fit as described above. We leveraged these non-linear normative curves, their confidence intervals, and first derivates separately for the left and right hemispheres and males and females, respectively. We modeled the 5th, 50th, and 95th centile values for each sex and hemisphere to visualize hemisphere-specific and sex-specific patterns of the claustrum volume change across the lifespan. Furthermore, developmental milestones of the claustrum (i.e., peaks of trajectories) were identified based on modeled medians (i.e., 50^th^ centiles) for each outcome measure (i.e., left and right claustrum as well as sex-stratified models).

Additionally, we used the 50th percentile (i.e., median) claustrum volumes to calculate sex- and hemisphere-related mean volumetric differences across the lifespan. The mean hemispheric asymmetry (%) was assessed using the formula: ((V_right_ – V_left_) / ((V_right_ + V_left_) / 2)) * 100 (Kong et al., 2020); where V_left_ and V_right_ represent the volume of the left and right claustrum respectively. Similarly, the sex-related mean volumetric differences (%) were calculated for each hemisphere using the formula: ((V_female_– V_male_) / ((V_female_ + V_male_) / 2)) * 100; where V_male_ and V_female_ represents the volume of the claustrum in males and females respectively.

##### Sex and hemisphere effect on claustrum volume

To analyze the specific effects of sex and hemisphere on claustrum volume while accounting for potential confounders such as age and TIV, we used an Ordinary Least Squares (OLS) regression model from the statsmodels package in Python (version 3.10.5). To account for potential scanner-related variability, TIV was first harmonized across sites using NeuroCombat (version 0.2.12, in Python) (Fortin et al., 2018), while preserving biological variation related to sex and age. The harmonized TIV values were then used as covariates in the NeuroCombat model to harmonize right, left, and averaged claustrum volumes, ensuring that site effects were minimized while preserving biological variation of sex and age. We constructed the following OLS models to systematically examine variable-specific effects: *Model 1: harmonized claustrum volume ∼ hemisphere + sex + age + harmonized TIV*, to assess the effect of hemisphere; *Model 2: right/left harmonized claustrum volume ∼ sex + age + harmonized TIV*, to evaluate the effect of sex separately for each hemisphere. The volumes derived from Model 1 and Model 2 are referred to as “TIV-adjusted claustrum volume” for consistency and clarity throughout the study. This terminology reflects the inclusion of TIV as a covariate in both models. We next aimed to estimate the effect of TIV on hemisphere and sex differences in claustrum volume; we repeated the above models without including TIV as a covariate. *Model 3: harmonized claustrum volume ∼ hemisphere + sex + age*, to quantify the influence of TIV on the estimated effects. The volumes derived from this model is called “absolute claustrum volume”. As an additional validation, we calculated claustrum volumes normalized by TIV by dividing the claustrum volume by each individual’s TIV. We then fitted Model 4, TIV-normalized claustrum volume ∼ hemisphere + sex + age, the result derived from this model is referred to as the TIV-normalized claustrum volume for throughout the study. (Supplementary 2.2).

## 3. Results

### 3.1. Segmentation model evaluation and segmentation quality

To assess the accuracy of automated claustrum segmentation, we calculated three performance metrics, namely VS, HD95, and DSC, on a selected data set of 40 subjects fairly distributed across the lifespan and sites, as defined in detail in the supplement methods part. The proposed automated segmentation method yielded a VS of 84.1%, an HD95 of 2.23 mm, and a DSC of 79.6% (Supplementary Figure S1, S2 Table S1). These results indicate high agreement with manual annotations, suggesting that the model provided an efficient and accurate automated segmentation of the claustrum. Detailed evaluation results are provided in the Supplementary material (Supplementary Figure S1, S2 and Table S1).

### 3.2. Modelling claustrum volumes across the lifespan

Centile normative values for averaged claustrum volume (Figure 3A), hemisphere-stratified (Figure 4 A, B), and sex-stratified (Figure 5 A1, A2, B1, B2) claustrum volumes are shown in respective figures. We evaluated all models by using metrics that quantify central tendency and distributional accuracy of each model (Fraza et al., 2021). More specifically, used metrics include (see Table 3): skew and excess kurtosis, which describe the estimated shape of the outcome Z-distribution and would ideally be close to zero; the explained variance (EV) describes the proportion of variance of the true value that is described by the model, with values being closer to 1 referring to a higher amount of variance in the data being explained by the model ; lastly, the mean standardized log loss, MSLL, favors simpler models, with more negative values indicating better model performance. Overall, as shown in Table 3, results for all metrics point to adequate model performance for all models (Rutherford et al., 2021). For example, skew, kurtosis, EV, and MSLL were 0.22, 1.22, 0.80, and 0.83, respectively, for the averaged claustrum volume model, indicating appropriate model predictions and performance. The specific claustrum volume trajectories will be described in more detail next.

**Table 3:**
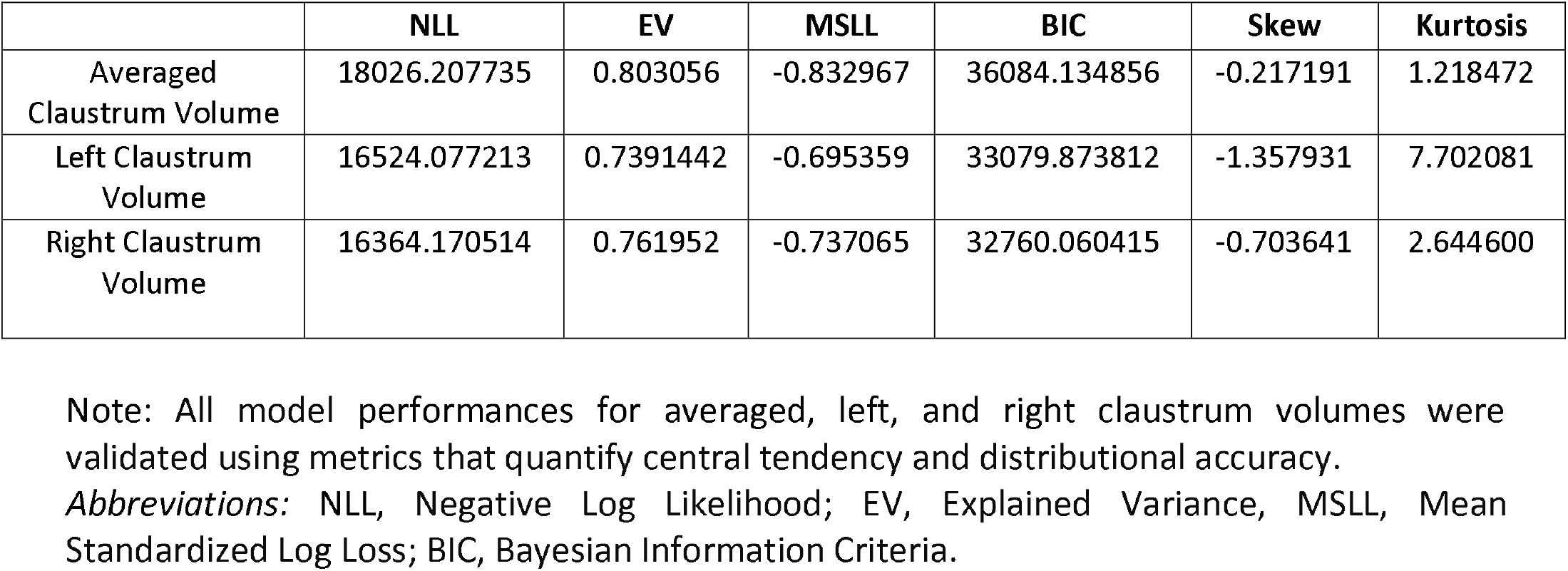
Error metrics for the evaluation of the normative claustrum volume models across the lifespan.

**Figure 3:**
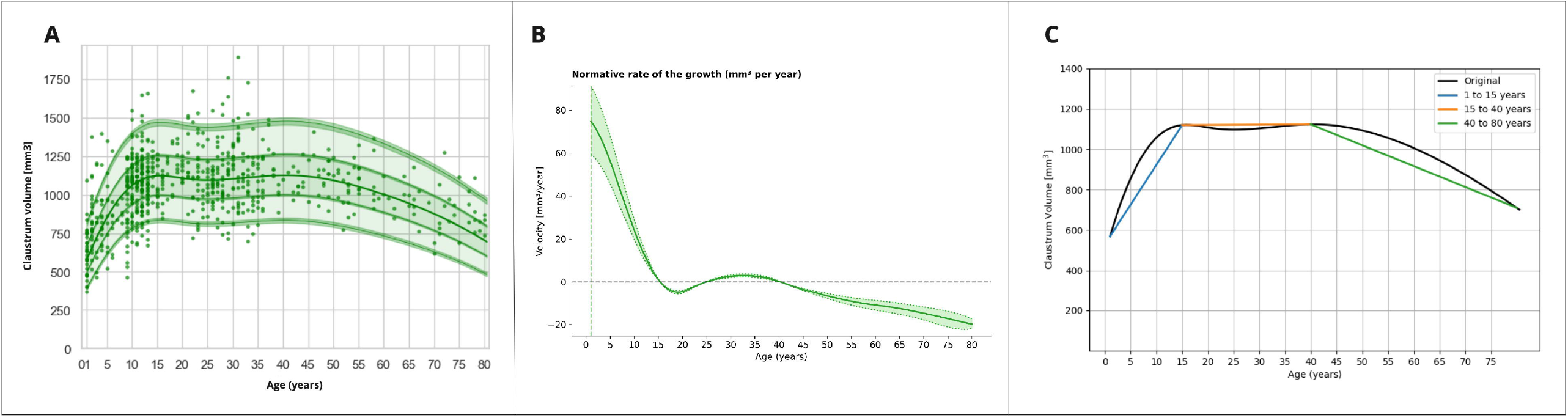
Averaged claustrum volume in humans: Normative life span trajectory and change rate for distinct age periods. A. Normative life span claustrum volume trajectory was estimated using BLR (site was used as a batch effect, and age, sex, and TIV were used as covariates). Claustrum volume was averaged across the left and right claustrum per subject. The 5^th^, 25^th^, 50^th^, 75^th^, and 95^th^ percentiles quantify the range of variation among individuals (dots). B. The rate of volumetric changes of the claustrum across the lifespan was estimated by the first derivatives of the median volumetric trajectories. C. Period-wise linear approximation of claustrum volume across the lifespan was applied to an increase phase (1–15 years, annual increase rate was ∼39.3 mm^3^/year.), a plateau phase (15–40 years), and a decrease phase (40–80 years, annual decrease rate was ∼10.325 mm^3^/year) to estimate the rate of change. *Abbreviations:* BLR: Bayesian Linear Regression

**Figure 4:**
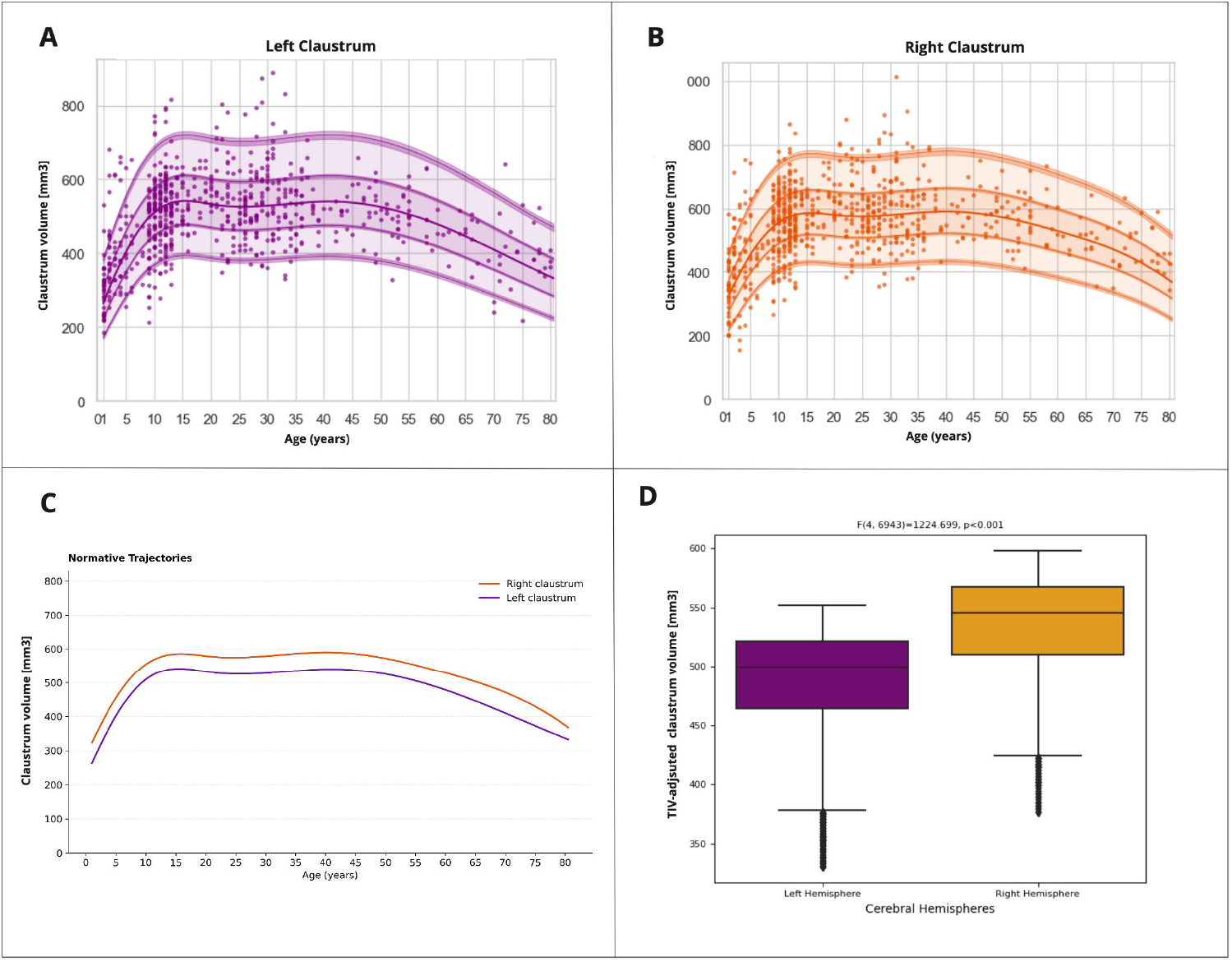
The effect of the hemisphere on claustrum volume. A-B: Normative life span trajectory of hemisphere-stratified claustrum volume. The 5^th^, 25^th^, 50^th^, 75^th^, and 95^th^ percentiles quantify the range of variation among lifespan-healthy individuals (black dots) in the right and left claustrum volume, respectively. C: Comparison of normative trajectories of the right and left claustrum volume across the lifespan. The median trajectories (50th centile values) were extracted for each hemisphere to visualize hemisphere-specific patterns of the claustrum volume change across the lifespan. D: Box plots show the TIV-adjusted right and left claustrum volumes derived from an OLS-based model adjusting for age, sex, and harmonized-TIV. Both left and right claustrum volumes were harmonized before analysis. The results show that the right claustrum volume is significantly higher than the left claustrum volume (p < 0.001).

**Figure 5:**
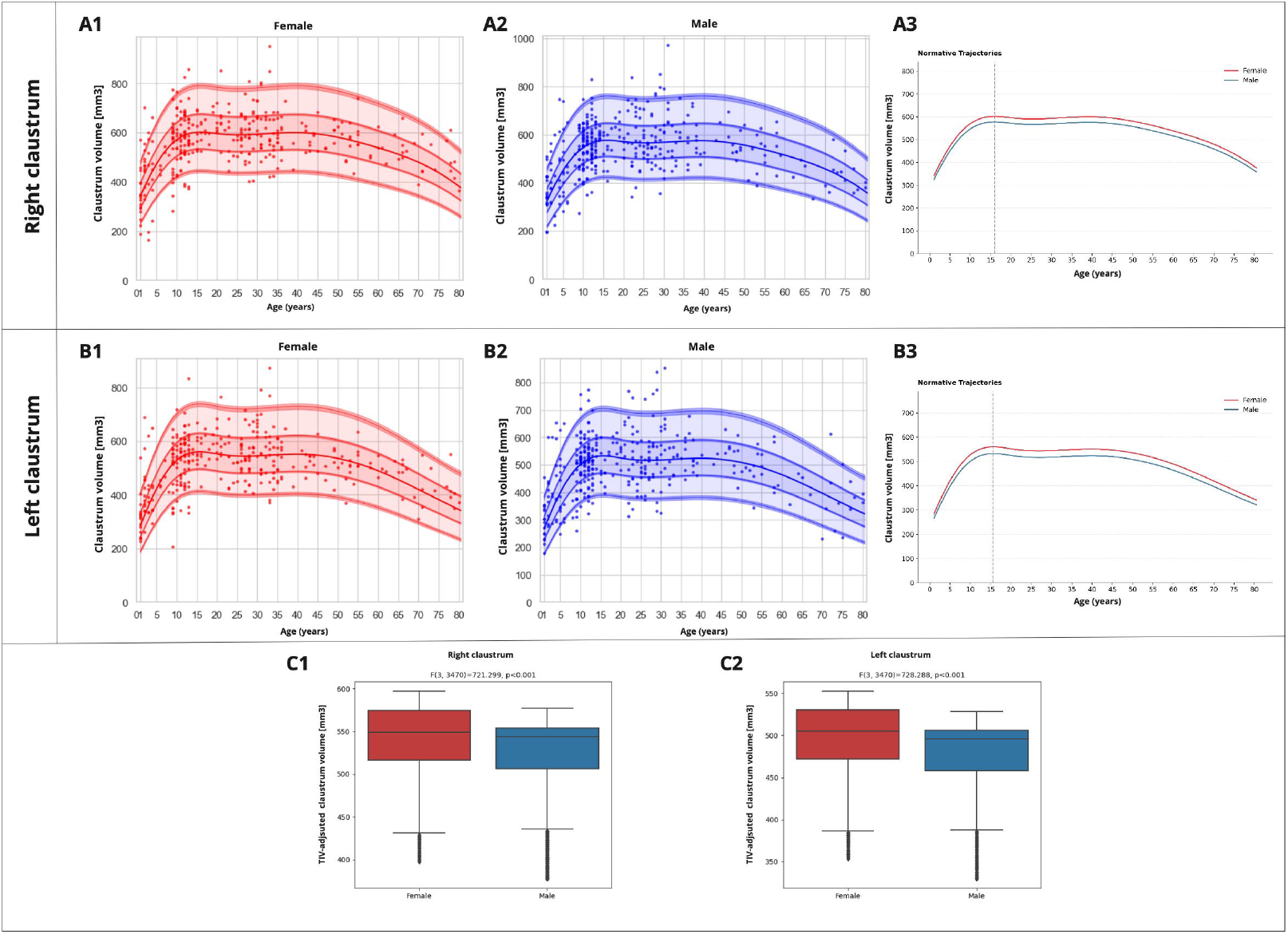
The effect of sex on the claustrum volume. Normative life span trajectory of sex-stratified claustrum volume. The 5^th^, 25^th^, 50^th^, 75^th^, and 95^th^ percentiles quantify the range of variation among lifespan-healthy individuals (black dots) in the right and left claustrum volume as a function of age (horizontal axis) and sex (female (A1, B1); male (A2, B2). Sex comparison of normative trajectories of the claustrum volume in females and males across the lifespan. The 50^th^ centile values were extracted for each sex to visualize sex-specific patterns of the claustrum volume change across the lifespan. The solid lines represent the median (50th percentile) trajectories. The vertical line indicates the age of maximum growth of each claustrum (A3, B3). The box plots display the predicted claustrum volumes for each region of interest (ROI), adjusted for sex, age, and harmonized-TIV. Box plots show the TIV-adjusted right and left claustrum volumes for females and males, derived from an OLS-based model adjusting for age, sex, and harmonized-TIV. Both left and right claustrum volumes were harmonized before the analysis. The results show higher claustrum volume in females than males for both hemispheres (p < 0.001) (C1, C2).

### 3.3. Lifespan trajectory of averaged claustrum volume and age effects

Lifespan curves showed that averaged claustrum volume increases from infancy to early adolescence, peaking in middle adolescence at about 15 years of age; then, claustrum volume is rather stable for a plateau phase, which lasts from early to middle adulthood of about 40 years of age; finally, volumes decline from middle adulthood to old age (Figure 3 A, B). Linear regression slopes were used to determine the rate of change based on non-linear trends in the median averaged claustrum volume. The findings indicate that the three age segments signify distinct phases for claustrum volume changes. The first phase, from 1 to 15 years, is characterized by a substantial increase, with an annual growth rate of approximately 39.300 mm^3^/year. The second phase, between 15 and 40 years, is identified as a plateau phase, with a slight decline in the annual change rate of approximately 0.153 mm^3^/year. The third phase, from 40 to 80 years, reflects a decrease in volume, with an annual decrease rate of about 10.325 mm^3^/year (Figure 3C). The annual rate of growth peaked between 1 and 2 years of age with approximately 83.002 mm^3^/year.

### 3.4. Hemisphere differences in claustrum volume across the lifespan

We optimized the BLR model specification and parameterization to estimate non-linear normative growth curves, their confidence intervals, and first derivatives separately for the claustrum in the right and left hemispheres. While the left and right claustrum volume trajectories were rather similar across the lifespan (Fig. 4A-B), detailed analysis revealed that the right claustrum is consistently larger than the left across all ages (the mean hemispheric asymmetry across all ages is 10.22% (SD = 2.42), Figure 4C). While the left claustrum volume peaked at age 15.7 years (95% bootstrap confidence interval (CI) 15.5–15.8), the right claustrum volume peaked at age 16.0 (95% bootstrap CI 15.8–16.1) (Table 4). When using a canonical OLS-based approach to further examine the effects of hemispheres on claustrum volume, accounting for the confounding variables age, sex, and harmonized TIV, we could confirm the finding of larger right claustrum volume: we found higher TIV-adjusted right claustrum volume compared to the TIV-adjusted left claustrum (β = 46.25, SE = 1.95, F(4, 6943) = 1225, p < 0.001) (Figure 4D), suggesting that on average, there is a difference in claustrum volume between left and right hemispheres in humans.

### 3.5. Sex differences in claustrum volume across the lifespan

Non-linear normative curves, their confidence intervals, and first derivates, separately for males and females for left and right claustrum, respectively, were then used to examine sex differences in claustrum volumes across the lifespan. We observed that right and left claustrum are consistently larger in females than in males across all ages (right claustrum, the mean difference across all ages is 4.27% (SD = 0.21); left claustrum, the mean difference across all ages is 5.37% (SD = 0.31), Figure 5 A3, B3). There was no difference in the peak age of claustrum volume between females and males, with both sexes showing peak volumes at 16.0 years for the right claustrum and 15.5 years for the left claustrum. The result of a canonical OLS-based approach on the effects of sex on claustrum volumes, accounting for confounding variables such as age and harmonized TIV, confirmed this finding by demonstrating that females have larger TIV-adjusted right claustrum (β = 20.39, SE = 3.38, F(3, 3470) = 721.299, p < 0.001) (Figure 5 C1). and TIV-adjusted left claustrum (β = 24.00, SE = 3.19, F(3, 3470) = 728.288, p < 0.001) compared to males (Figure 5 C2). These findings suggest that sex has an influence on relative claustrum volume in humans, with relatively larger volumes in females. We confirmed this finding by further control analyses in the Supplement (Supplementary Figure S4): while absolute claustrum volumes of males were larger than those of females (β = 40.59, SE = 3.27, F(3, 3471) = 397.858, p < 0.001), TIV-normalized claustrum volumes (i.e., claustrum volume divided by TIV) were larger in females across the lifespan (β = 32.78, SE = 3.08, F(2, 3471) = 420.484, p < 0.001).

## 4. Discussion

Using T1w-MRIs of 3,474 healthy individuals from 1 to 80 years of age and automated claustrum segmentation, we observed hemisphere- and sex-distinct claustrum volume lifespan trajectories. In general, claustrum volumes increase from 1-15 years, are stable up to 40 years, and then decrease up to 80 years. The right claustrum is larger than the left one. Female TIV-adjusted claustrum volumes are larger than in males. To the best of our knowledge, this is the first large sample study of claustrum structure across the lifespan in humans, demonstrating age-, hemisphere-, and sex-sensitive claustrum trajectories. This result provides a comprehensive framework for claustrum lifespan trajectories relevant for studying neurodevelopmental and neurodegenerative effects on the claustrum.

### 4.1. Lifespan trajectories of claustrum volume

We found a distinct association between age and claustrum volume across the lifespan. Claustrum volumes increase constantly during childhood, reaching its peak around 15–16 years of age. This period of increase is followed by a period of stability lasting until the end of the third decade of life, with a subsequent period of decline during the second half of middle adulthood and late adulthood. This pattern of claustrum volume change aligns broadly with the inverted U-shaped growth curves observed in thalamus, amygdala nuclei and the hippocampi(Coupé, Catheline, Lanuza, & Manjón, 2017; Dima et al., 2022). But, while these structures typically peak later in life (generally within the first 2–3 decades) (Coupé et al., 2017; Dima et al., 2022), the peak of claustrum volume development more closely resembles the developmental pattern of the basal ganglia (striatum, pallidum), which also reach their maximum volume during adolescence (approximately at 14 years) (Bethlehem et al., 2022b; Narvacan, Treit, Camicioli, Martin, & Beaulieu, 2017).

However, comparing claustrum growth rates with those of other subcortical structures remains challenging, for example due to methodological variations in peak age estimations across studies. These discrepancies stem not only from differences in the total age range analyzed but also from variations in the starting age of the samples, as large volumetric increases occur in early postnatal life (<2 years) (Knickmeyer et al., 2008; Narvacan et al., 2017). In line with this, our findings indicate that largest claustrum volume changes per year occur between 1 and 2 years of age, a dynamic and critical stage in postnatal brain development. This rapid early growth suggests that the claustrum may play an important role in cognitive development as well as the potential pathogenesis of neurodevelopmental disorders.

Supporting this notion, Neubauer et al. (2023) reported that preterm neonates exhibit increased claustrum volume compared to term infants, alongside alterations in microstructure. However, Hedderich et al. (2021) found that while claustrum volume in adults born preterm were not significantly different compared to controls, microstructural differences exist. Furthermore, a lifespan study on autism by Wegiel et al. (2014) revealed significant developmental alterations in the claustrum across different life stages. Their findings indicate that the most pronounced neuronal soma deficits occur in children with autism (4–8 years old), compared to teenagers and adults (Wegiel et al., 2014). These observations align with the hypothesis that early-life deviations in claustrum macrostructure may contribute to atypical trajectories in neurodevelopmental disorders.

### 4.2. Hemisphere differences in claustrum volume across the lifespan

In addition, our findings reveal a consistent hemisphere difference in claustrum volumes, with the right claustrum being larger than the left across all ages. This result is independent of TIV influences, as the analysis of left-right claustrum differences did not change when including TIV as a covariate of no interest.

The observed volume differences between the right and left claustrum align with findings in adults, although the degree of asymmetry varies across studies (Berman, Schurr, Atlan, Citri, & Mezer, 2020; Milardi et al., 2015; Torgerson et al., 2015). The claustrum exhibits hemispherical asymmetry, with the right claustrum showing a slightly larger average absolute volume (829 mm^3^) compared to the left (706 mm^3^) (Kapakin, 2011; Nikolenko et al., 2021). This rightward asymmetry parallels hemispherical asymmetry patterns observed in other cortical structures such as posterior cortex, including the temporal lobe (lateral and medial parts), and the medial occipital cortex (Kong et al., 2018; Roe et al., 2023), hippocampus and subcortical structures such as amygdala, nucleus accumbens and caudate nucleus (Guadalupe et al., 2017; Kong et al., 2022; Ocklenburg, Peterburs, & Mundorf, 2022; Okada et al., 2016) which have also been shown to exhibit rightward volumetric dominance.

Hemispherical asymmetry in brain structure volume reflects distinct patterns in structural design that are linked to specific functions (Kuo & Massoud, 2022). The functional implications of the hemispheric differences in claustrum volume remain unclear; however, Rodriquez et al. provided evidence of lateralized functional connectivity in the resting-state claustrum in healthy adults. They demonstrated that the right claustrum exhibits stronger connectivity with the frontoparietal and dorsal attention networks, supporting the hypothesis of claustral asymmetry. (Rodríguez-Vidal, Alcauter, & Barrios, 2019). Additionally, previous studies suggest that the right claustrum may be more actively involved in certain cognitive processes. For instance, greater right claustrum activation was reported during modal sensory integration of conceptually related objects (Naghavi, Eriksson, Larsson, & Nyberg, 2007) and suppressing natural urges (Lerner et al., 2009). Based on these findings, we propose that the larger claustrum volume in the right hemisphere may be linked to its more prominent role in sensory integration and attentional control of the claustrum across the lifespan.

In addition to the asymmetrical macrostructure of the claustrum in healthy populations, asymmetrical involvement of the claustrum was observed in the severity of certain conditions in Alzheimer’s disease and schizophrenia (Bruen et al., 2008; Cascella et al., 2011). Notably, significant inverse correlations between ratings of the severity of delusions and volumes of the left claustrum were observed in Alzheimer’s disease (Bruen et al., 2008) and schizophrenia (Cascella et al., 2011). The need for further investigation is whether the claustrum volumetric difference between hemispheres is functionally linked to lateralized cognitive functions or whether it contributes to vulnerability to neuropsychiatric conditions that exhibit hemispheric lateralization (e.g., schizophrenia).

### 4.3. Sex differences in claustrum volume across the lifespan

Finally, we observed that the TIV-adjusted volumes of the right and left claustrum are consistently larger in females than in males across all age groups. Additionally, TIV significantly influenced claustrum volume, as evidenced by the opposite-sex effect on absolute claustrum volume, with males exhibiting relatively larger volumes in both hemispheres throughout the lifespan.

Previous studies have reported that, on average, males tend to have a larger absolute claustrum volume compared to females (Baizer, Sherwood, Noonan, & Hof, 2014; Ruigrok et al., 2014). While our findings are in line with the sex differences in absolute claustrum volume, we found that females have larger claustrum volume when including TIV as a covariate. This pattern is consistent with previous observations in other cortical and subcortical structures. Males generally exhibit larger total brain volumes and greater variability in brain structure compared to females across the lifespan. (Diáz-Caneja et al., 2021). However, when controlling for TIV, females show greater grey matter volume than males in several cortical regions, including the prefrontal and superior parietal cortices, as well as the superior temporal sulcus, orbitofrontal cortex, and posterior insula (S. Liu, Seidlitz, Blumenthal, Clasen, & Raznahan, 2020; Lotze et al., 2019). Similarly, the caudate nucleus has been found to be relatively larger in females (Giedd, Castellanos, Rajapakse, Vaituzis, & Rapoport, 1997). Notably, studies on limbic structures such as the hippocampus (Tan, Ma, Vira, Marwha, & Eliot, 2016) and amygdala (Marwha, Halari, & Eliot, 2017) have reported that while their absolute volume was larger in males, this difference disappears when TIV is accounted for. TIV is a key global morphometric measure with the greatest significant effect on regional subcortical volumes and cortical surface area measures (Ge et al., 2024). Our findings suggest that TIV plays a crucial role in explaining sex differences in claustrum volume, similar to its influence on other specific brain structures. Specifically, we propose that the relatively higher claustrum volumes in females exceeds the proportional increase in TIV observed in males.

Although the underlying causes of this sex difference in the claustrum remain unclear, sex-based variations in brain structure are commonly attributed to genetic, hormonal, and environmental influences (Diáz-Caneja et al., 2021; Ruigrok et al., 2014). The observed sex differences in claustrum volume may have implications for understanding sex-specific vulnerabilities to neurological and psychiatric conditions, as well as differences in cognitive and emotional processing.

Altogether, our findings highlight significant hemisphere and sex differences in claustrum volume across the lifespan. The demonstrated hemisphere and sex differences in claustrum volume require a more nuanced approach in neuroimaging research, particularly in studies examining the claustrum’s role in cognitive and emotional processing. Future research should explore the functional correlates of these structural differences, potentially employing multimodal imaging techniques to elucidate the relationship between claustrum morphology and neural activity patterns.

### 4.4. Strengths and Limitations

The unique strengths of our study are the availability of a large and diverse sample size and the use of deep learning algorithms for claustrum segmentation. We trained a model on a lifespan cohort, and the model achieved satisfactory segmentation results across large sample size ranging from 1 to 80 years old. Furthermore, the BLR modeling approach allowed for harmonization across heterogeneous data sets, enabling the generation of normative, hemisphere- and sex-stratified claustrum growth charts by incorporating individual variation. The application of BLR to model normative trajectories provides a robust framework for understanding individual differences in claustrum volume across demographic categories. Moreover, we supported our normative modeling analysis with canonical OLS regression analysis to investigate the sex and hemisphere differences in claustrum volume. We believe that the established trajectories of claustrum volume across the lifespan by combining different statistical approaches provide valuable insight for future studies investigating the role of the claustrum in various neurodevelopmental and neurodegenerative disorders.

There are some limitations to this study. First, participants younger than one year were not included. We believe that accurately capturing claustrum volume dynamics in this early period would require a significantly larger sample size and diving more specific into small age range groups due to the rapid and highly dynamic changes in brain development (Knickmeyer et al., 2008). Second, the use of multiple datasets with different meta information made it challenging to systematically assess the correlation between claustrum volume and cognitive or attentional performance across the lifespan.

## 5. Conclusion

In summary, this study provides valuable insights into the development of claustrum volume across the lifespan, highlighting significant age, hemisphere, and sex differences. By elucidating these biological factors, we contribute to a deeper understanding of structural characteristics, growth pattern, and potential implications of claustrum volume for neurodevelopmental and neuropsychiatric research.

## CRediT authorship contribution statement

**Sevilay Ayyildiz:** Writing – review & editing, Writing – original draft, Data curation, Visualization, Methodology, Investigation, Formal analysis, Conceptualization. **Antonia Neubauer:** Writing – review & editing, Methodology, Investigation. **Melissa Thalhammer:** Writing – review & editing, Validation, Methodology, Conceptualization. **Hongwei Bran Li:** Writing – review & editing, Methodology, Investigation. **Jil Wendt**: Writing – review & editing, Data curation, Conceptualization. **Aurore Menegaux:** Writing – review & editing, Data curation. **Rebecca Hippen:** Writing – review & editing, Conceptualization. **Benita Schmitz-Koep:** Writing – review & editing, Conceptualization. **David Schinz:** Writing – review & editing, Conceptualization. **Claus Zimmer:** Writing – review & editing, Resources. **Behcet Ayyildiz:** Writing – review & editing, Data curation, Conceptualization.**Abdullah Ors:** Writing – review & editing. **Belgin Bamac:** Writing – review & editing, Project administration, Supervision.

**Dennis Hedderich:** Writing – review & editing, Project administration.**Christian Sorg:** Writing – review & editing, Supervision, Resources, Project administration, Conceptualization.

## Acknowledgements and Disclosure

Data was provided by the public datasets, including the Lifespan Baby Connectome Project, the Calgary Preschool MRI Dataset, the Adolescent Brain Cognitive Development, and the Human Connectome Project Lifespan. We sincerely thank data providers and the participants whose contributions made this research possible.

## Data Availability Statement

The data used in this study come from the UNC/UMN Lifespan Baby Connectome Project (BCP), the Calgary Preschool MRI Dataset (CPD) (https://osf.io/axz5r/), a selected dataset from the Adolescent Brain Cognitive Development (ABCD) study, 5.1 release, primarily including participants aged between 9 and 12 years (https://abcdstudy.org), the Human Connectome Project (HCP) Development dataset, the HCP Young Adult dataset, and the HCP Aging dataset (https://www.humanconnectome.org), which are freely available after appropriate data usage agreements.

The code for automated claustrum segmentation in neonatal brain MRI is available on GitHub: https://github.com/hongweilibran/claustrum_multi_view.

