## Supplementary Material for "Claustrum volume in humans – lifespan trajectory and effect of age, hemisphere, and sex"

The claustrum in humans – across the life span, inverted-U-like volume changes with relatively larger volumes in the right hemisphere and females

by Sevilay Ayyildiz et al.

### 1  Supplementary Methods:

#### 1.1 Claustrum segmentation

Our study encompasses a comprehensive approach on automated deep-learning-based claustrum segmentation across the lifespan. The segmentation modeling process entails several steps, including data preparation, augmentation, network architecture definition, and training with meticulously selected hyperparameters. First, the data preparation steps were carried out as described in the main paper. Then, manual claustrum segmentation was performed for selected subjects to create a segmentation model.

##### Manual segmentation

50 subjects were randomly selected from across all datasets, ensuring representation across various age groups and sites, for manual segmentation of the claustrum by an experienced neuroanatomist (S.A.). Manual annotations of the claustrum followed a modified manual claustrum segmentation protocol adapted from Davis (2008) (Davis, 2008; see also Hedderich et al., 2021 for more details) using ITK-SNAP-v 4.0.2 on a Wacom Intuos M tablet (Wacom, Kazo, Saitama, Japan) (Yushkevich et al., 2006).

Subsequently, these scans were partitioned into a training set of 40 subjects and a test set comprising 10 scans for evaluation. The remaining 3498 scans did not undergo manual segmentation.

##### Image augmentation

Various data augmentation techniques, including scaling, shifting, rotation, and shearing, were employed to augment the diversity of training samples, thereby enhancing the model's generalization capability. Additionally, intensity augmentation was performed to enhance the model's robustness against variations in image intensity (Neubauer et al., 2022).

##### Segmentation algorithm: U-net convolutional neural network architecture and transfer learning

We applied a deep learning-based algorithm, whose models and code are publicly available (https://github.com/hongweilibran/claustrum_multi_view). A supervised deep-learning approach originally developed for adults (Li et al., 2021) is adapted for automated lifespan claustrum segmentation using transfer learning (Neubauer et al., 2022). The architecture utilizes the U-Net convolutional neural network, comprising a down-convolutional segment for feature extraction from T1w input scans and an up-convolutional segment to finally categorize each pixel as claustrum or non-claustrum. Skip connections between these segments are integrated to facilitate seamless information flow between the contracting and expanding pathways. The network architecture is described in detail in Li et al. (Li et al., 2021). To optimize performance on lifespan data, we performed transfer learning by initializing the model with weights from a pre-trained adult claustrum segmentation network and fine-tuning it on lifespan MRI scans (Neubauer et al., 2022).

##### Model training and hyperparameters

The training process is guided by the Dice coefficient loss function, which quantifies the similarity between predicted and ground truth manual segmentations. Coronal and axial deep convolutional neural networks are trained on 2D single-view slices obtained after parsing 3D MRI volumes into axial and coronal views, a strategy inspired by the advantageous multi-view approach proposed in prior literature (Li et al., 2021; Neubauer et al., 2022). During testing, predictions are automatically combined at the voxel level (Li et al., 2021).

Critical hyperparameters such as batch size, learning rate, and number of epochs are pre-defined to optimize training efficacy. Each model undergoes training for 30 epochs to mitigate the risk of overfitting (Neubauer et al., 2022). The batch size of 60 dictates the number of samples processed in each iteration, while the learning rate of 0.0002 regulates the magnitude of optimization steps.

##### Model evaluation

We trained and aggregated three axial and coronal view models to enhance the model robustness, culminating in a unified model at the voxel level. We assessed the segmentation performance using three different evaluation metrics (Li et al., 2021; Neubauer et al., 2022). Volumetric similarity (VS) is defined as the similarity between the volumes of the claustrum manual segmentation mask and the predicted segmentation mask. Values closer to 1 (or 100%) refer to better overlap between the predicted and manual segmentation volumes. 95th Percentile of the Hausdorff Distance (HD95) is a score to measure surface distance between manual segmentation and predicted mask. A lower HD95 value (close to 0) means better accuracy in surface alignment. The Dice similarity coefficient (DSC) quantifies the spatial overlap between manual segmentation and predicted mask. Values closer to 1 (or 100%) indicate to better overlap between the manual and predicted segmentations.

To assess the performance of the axial and coronal multi-view technique on the training/validation set, a stratified 5-fold cross-validation was conducted. Each fold comprises an 80/20 split, with 80% of scans allocated to the training set and the remaining 20% to validation. Following five iterations, all subjects undergo evaluation in the validation phase. The model was thus optimized on 40 subjects. The combined model was evaluated on a test set of 10 subjects. Then, the combined model is applied to the remaining 3498 subjects’ scans.

### 2 Supplementary Results:

#### 2.1 Segmentation accuracy

To evaluate the segmentation performance, we calculated the volumetric similarity (VS), Hausdorff distance (95th percentile) (HD95), and Dice similarity coefficient metrics to assess various aspects, as delineated in prior studies (Li et al., 2021; Neubauer et al., 2022). The results of the evaluation process for claustrum segmentation are shown in **Figure S1 and S2 and Table S1**.

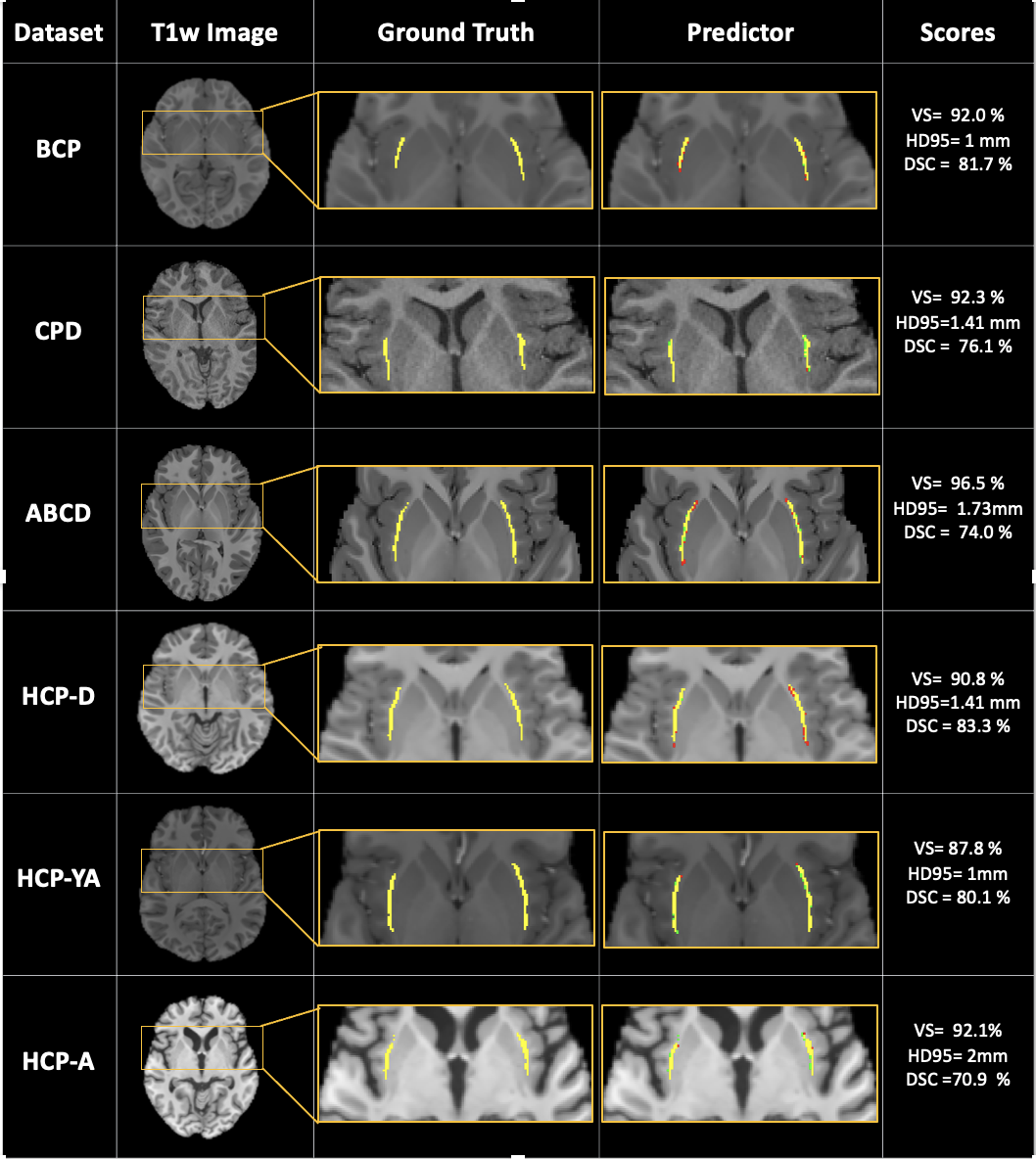

Figure S1: Segmentation of six example cases, one per dataset.

This figure presents manual (ground truth) and automatic (predicted) segmentation results for six example cases, one from each dataset. The predicted segmentation masks are compared to the ground truth to evaluate combined model performance. In the predicted segmentation masks, the yellow pixels represent true positives, the green ones represent false negatives, and the red ones represent false positives. The accuracy of each example is reported using the Dice Similarity Coefficient (DSC), Volumetric Similarity (VS) and the 95th percentile of Hausdorff Distance (HD95). The values are reported as mean in the example sample cases. Higher VS and DSC values indicate better segmentation performance, while lower HD95 values means better accuracy in surface alignment.

*Abbreviations:* BCP, UNC/UMN Lifespan Baby Connectome Project; CPD, Calgary Preschool MRI Dataset; ABCD, Adolescent Brain Cognitive Development; HCP, Human Connectome Project; HCP-D, HCP Development; HCP-YA, HCP Young Adult; HCP-A, HCP Aging.

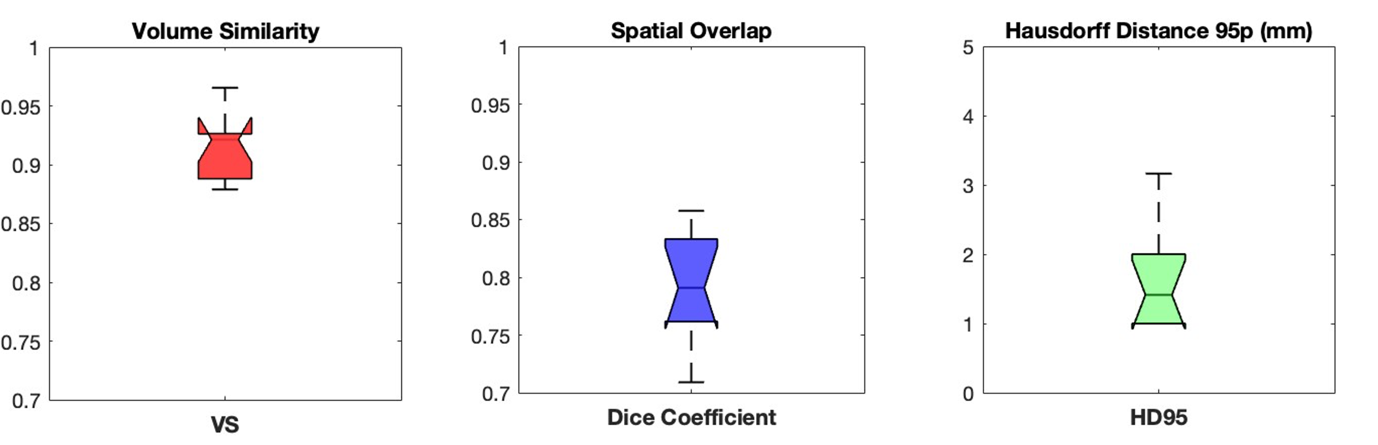

**(%)**

**(%)**

Figure S2: Model performance of automated claustrum segmentation.

Segmentation performance of the proposed method on the test set (automated segmentation). The segmentation accuracy of the automated segmentation was evaluated using three standard performance metrics: Volumetric Similarity (VS), 95th percentile of the Hausdorff Distance (HD95), and Dice Similarity Coefficient (DSC). These metrics assess volumetric agreement, boundary accuracy, and spatial overlap, respectively. The corresponding values for these accuracy metrics are reported in Table S1.

Table S1: Model performance of automated claustrum segmentation.

| VS (%)  Median, [IQR] | DSC (%)  Median, [IQR] | HD95 (%)  Median, [IQR] |
| --- | --- | --- |
| 84.1, [88.7, 93.2] | **79.6, [77.3, 83.4]** | **2.23, [1.41, 2.82]** |

Note: Segmentation performance of the proposed method on the test set (automated segmentation). This table presents the segmentation accuracy of the proposed method based on three performance metrics: Volumetric Similarity (VS), 95th percentile of the Hausdorff Distance (HD95), and Dice Similarity Coefficient (DSC). The values are reported as mean ± standard deviation across the test set. Higher VS and DSC values indicate better segmentation performance, while lower HD95 values suggest improved boundary agreement.

#### 2.2 Impact of the hemispheres and sex on claustrum volume across the lifespan: control analyses for absolute and TIV-normalized claustrum volume

To further investigate the impact of hemisphere and sex on claustrum volumes, we conducted several control analyses using Ordinary Least Squares (OLS) regression, mainly to account for the impact on TIV claustrum volume differences between hemispheres or sexes, respectively. Harmonized claustrum volumes and TIV values were used in all analyses.

##### 2.2.1 Control analysis for the impact of the hemisphere on claustrum volumes

In the first control analyses, in order to assess whether claustrum volume differences between hemispheres are influenced by TIV, we applied, first, TIV metric normalization by dividing the right and left claustrum volumes by TIV. The model (TIV-normalized claustrum volume ~ hemisphere + sex + age) was used. The claustrum volume in this analysis is referred to as “*TIV-normalized claustrum volume*”. The results showed that the right claustrum volume is significantly higher than the left in the TIV-normalized measure (p < 0.001) (Figure S3A).

In the second control analysis, the model (claustrum volume ~ hemisphere + sex + age) was fitted to analyze the effect of the hemisphere on claustrum volume without accounting for potential confounders TIV. The claustrum volume in this analysis is referred to as “*Absolute claustrum volume*”. Our findings suggest that the left claustrum volume is significantly higher than the right in the absolute volume measure (p < 0.001) (Figure S3B).

Taken together, both control analyses confirm larger right claustrum volume across ages, independent from TIV differences.

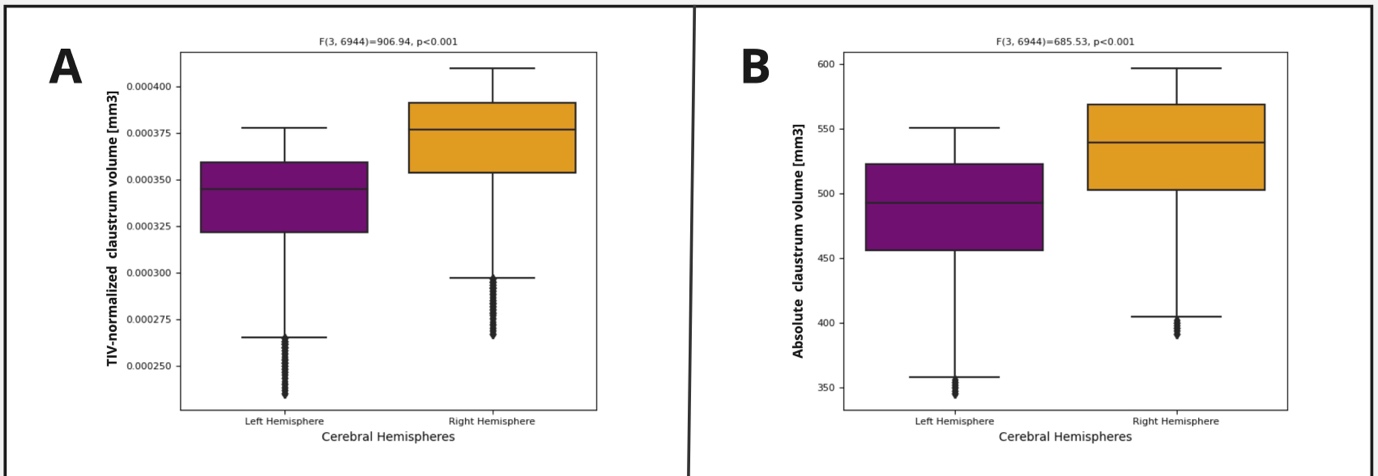

###### Supplementary Figure S3: Impact of the hemisphere on the claustrum volume, with and without accounting for TIV. An OLS-based model (with sex and age as covariates) was used to assess differences in the left and right claustrum volumes. (A) TIV-normalized claustrum volume (B) Absolute claustrum volume. The right claustrum volume is significantly higher than the left one for both the TIV-normalized measure (p < 0.001) and the absolute volume measure.

##### 2.2.2 Control analysis for the impact of sex on claustrum volumes, accounting for differences in TIV across sexes.

In order to assess whether claustrum volume differences between sexes are influenced by TIV, we applied, first, TIV metric normalization by dividing the right and left claustrum volumes by TIV. The model (TIV-normalized averaged/right/left claustrum volume ~ sex + age) was used. Such *TIV-normalized claustrum volume* was larger in females than in males (p < 0.001) (Figure S4A).

Then we performed the same analysis but using absolute claustrum volumes instead of TIV adjusted volumes. Based on the model (averaged/right/left claustrum volume ~ sex + age), we observed that the absolute claustrum volume in males is significantly higher than that of females (p < 0.001) (Figure S4B).

Taken together, these findings demonstrate that TIV-relative but not absolute claustrum volumes were larger in females than in males.

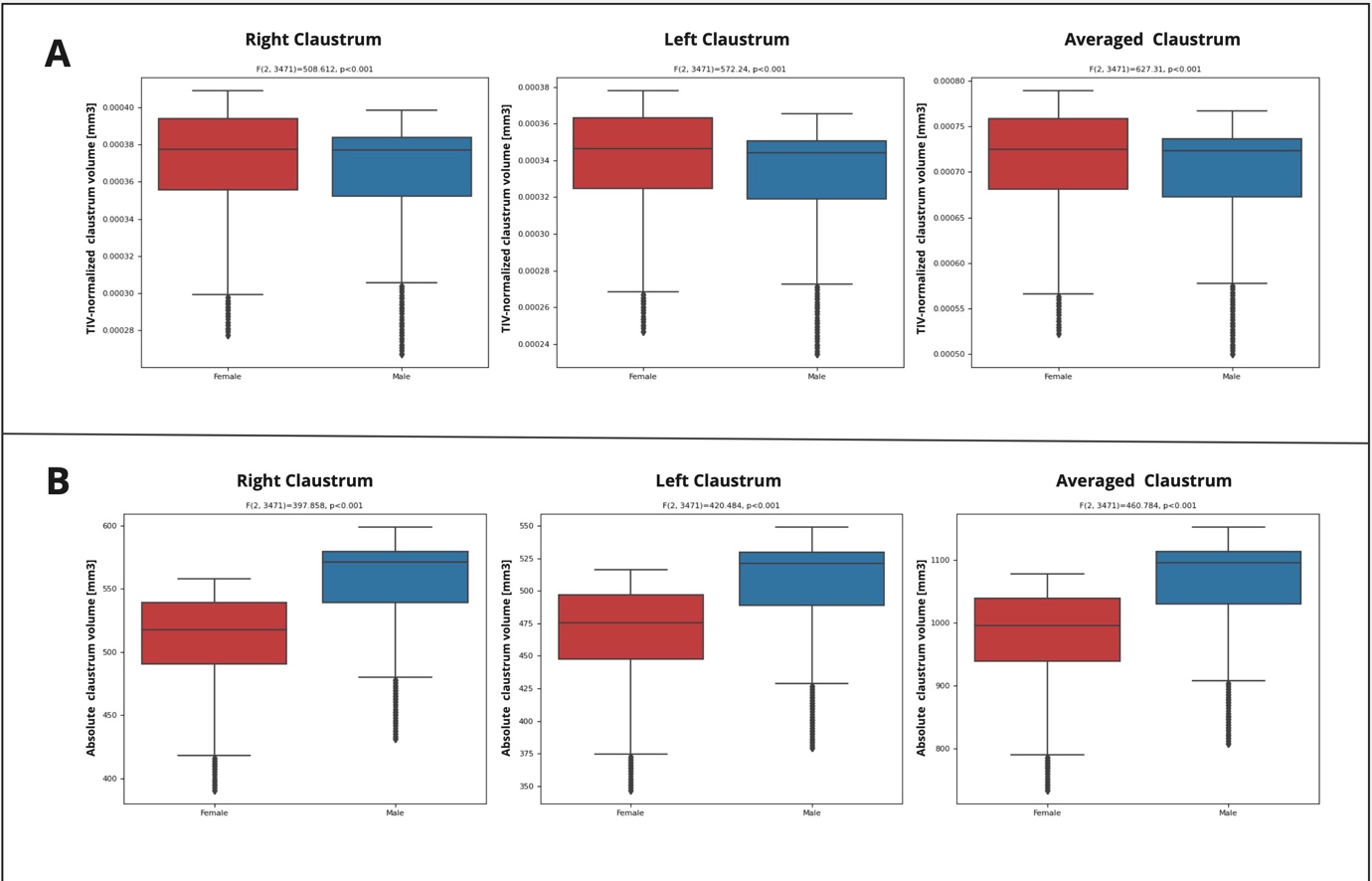

Supplementary Figure S4: Impact of sex on the claustrum volume.

An OLS-based model (with age as a covariate) was used to assess sex differences in the left, right, and averaged claustrum volumes, with and without accounting for differences in TIV. (A) TIV-normalized claustrum volume (B) Absolute claustrum volume. The TIV-normalized claustrum volume in females is significantly larger than that in males (p < 0.001), while the absolute claustrum volume is larger in males (p<0.001).
